# Opportunistic diver-assisted eDNA sampling unpicks fine-scale ecological and conservation signals in tropical reef fishes

**DOI:** 10.1101/2025.06.11.659134

**Authors:** Giulia Maiello, Victor Burgos Rubio, Lara Allen, Erika F. Neave, Rodney Bonne, Patsy Theresine, Stefano Mariani

## Abstract

Coral reefs are emblematic ecosystems of astonishing biological diversity and significant economic value for many countries, yet they are increasingly vulnerable to direct and indirect human-related threats, such as heavy tourism and global warming. To assess the biodiversity status, track assemblage changes and forecast future trends of these megadiverse ecosystems, there is pressing need for accurate, rapid and comprehensive assessments of fish communities. Here, we employed a versatile low-tech passive environmental DNA collector, the ‘metaprobe’, in association with the recreational activity of scuba divers to collect eDNA data from reefs in the Seychelles archipelago and assess both the taxonomic and functional fish diversity of the area. Using a fish-specific 12S marker on eDNA metaprobe samples collected during 18 dives, we detected 174 fish taxa, eight elasmobranchs and 166 teleosts, corresponding to 112 unique functional entities and including iconic tropical reef taxa, endangered elasmobranchs and commercially valuable species. Despite the geographic proximity of dive sites, we identified ecological patterns of fish composition and community structure influenced by both the presence of marine protected areas (MPAs) and the type of predominant substrate (i.e., coral vs rocky). Ecological differences between MPA/non-MPA and coral/rocky habitats involved both taxonomic and functional community elements, which returned very similar patterns of significantly higher fish diversity in protected and coral areas. We argue that simple, inexpensive eDNA-based collection tools in association with recreational activities, such as scuba diving, provide a promising approach for upscaling ecological data collection and expanding biodiversity monitoring, while engaging and empowering the public.

## 1. Introduction

Coral reefs are the main marine biodiversity hotspot of the planet (Fisher et al., 2015); despite covering less than 1% of the world’s surface, about one-fourth of all known marine species feed, hide and breed on and around them. Coral ecosystems are presently facing serious threats (Williams et al., 2019); their rapid degradation is mainly driven by global warming, land-based pollution and human activities such as high fishing pressure, often due to poor regulation or even illegal activity (Hughes et al., 2018; McClure et al., 2021; Newton et al., 2007). All of these stressors together are increasing the vulnerability of coral reef fishes, leading to declines in abundance and even local extinction of some species (Graham et al., 2011). Coral reefs also constitute major tourist attractions and often represent a vital source of income in many countries (Spalding et al., 2017). There are consequently considerable incentives to understand coral communities and monitor their changes, towards appropriate conservation plans and sustainable use of biological resources (Cinner et al., 2020)

Monitoring megadiverse ecosystems such as coral reefs, however, presents huge challenges (Plaisance et al., 2011). Traditionally, marine species inventories have been produced using various approaches, such as direct capture of organisms, underwater visual census (UVC) and/or using remote optical/acoustic methods (Alter et al., 2022; Evans et al., 2017; Stat et al., 2019; Thomsen et al., 2012), each with several limitations. Capture-based methods are invasive and environmentally impactful, and all of these methods require high taxonomic expertise, are time-consuming, and considerably depend on weather conditions, limiting both the sample size and the spatial and temporal coverage (Andruszkiewicz et al., 2017; Sigsgaard et al., 2017). Environmental DNA (eDNA) has proven its worth as an effective marine bioassessment approach; its universality, low-impact, and relative simplicity, have opened new opportunities for upscaling data collection from the ocean (Gilbey et al., 2021), and its success appears evident even in highly diverse coral reef ecosystems (Nguyen et al., 2020; Polanco Fernández et al., 2021; Sigsgaard et al., 2019; West et al., 2020).

Beyond the recording of taxonomic diversity, which describes the different taxonomic units, without considering ecological differences among species (Bosch et al., 2017), a more robust understanding of ecosystems structure and its changes in response to natural and anthropogenic perturbations comes from the characterisation of functional diversity, that is, the diversity in functional traits across all the species in the assemblage (D’Agata et al., 2014; McGill et al., 2006). Trait-based metrics have thus become more common in monitoring the status of fish communities in various ecosystems (Aune et al., 2018; Rincón-Díaz et al., 2018; Toussaint et al., 2016). Recent applications demonstrated that marine eDNA data can provide reliable insights into the functional diversity of fish communities (Condachou et al., 2023; Marques et al., 2021; Rey et al., 2023; Roblet et al., 2024). Aglieri et al. (2021) even found that eDNA metabarcoding outperforms traditional survey methods (such as visual census, underwater video and small-scale fishery) in the ability to investigate functional diversity, because of the DNA molecules’ inherent lesser bias towards any functional category and its consequently greater ability to efficiently capture a wider spectrum of traits.

Collecting marine eDNA, however, does not come without its own challenges, especially the need for laborious, time-consuming, water-filtration procedures, which significantly limit eDNA sampling effort (Bessey et al., 2021; Hinlo et al., 2017) and prevent a broader application of this approach especially in remote and non-equipped locations (Hansen et al., 2020). Such limitations have spurred growing efforts towards the development of passive methods to collect eDNA from the water (Bessey et al., 2021; Jeunen et al., 2022, 2024; Verdier et al., 2021), one of the most versatile of which is the ‘metaprobe’: a reusable custom-made 3D-printed sphere, that passively collects eDNA through rolls of gauze placed inside it (Maiello et al., 2022; 2023). The metaprobe represents an agile, low-tech and inexpensive sampling tool, offering a less labour-demanding alternative to water filtration, and can easily be operated by everyone, in the most disparate environmental settings.

Here we opportunistically combined the metaprobe with the recreational activity of scuba divers to collect eDNA samples in 18 locations in the North-Western coast of Mahé, the largest island of the Seychelles archipelago, in the Indian Ocean. The sampled area encompasses two marine protected areas (i.e., Baie Ternay and Port Launay Marine National Parks) and unprotected areas, as well as different types of marine substrates, primarily coral reefs and granitic rocks. We used metaprobe-based environmental DNA data to assess the taxonomic and the functional fish diversity of this hyperdiverse tropical area, specifically aiming at: (i) evaluating the benefits of attaching metaprobes to recreational scuba divers to capture the high fish diversity of the area, with an emphasis on threatened and commercially valuable species; (ii) evaluating the degree of influence of the type of substrate (i.e., coral or granitic) and the presence of a marine protected area on the taxonomic and functional richness and composition of fish communities.

## 2. Material and methods

### 2.1. Sample Collection

Sample collection took place during the spring/summer season of 2023 from 18 sites along the western coast of Mahé, Seychelles, in the Indian Ocean (Fig. 1A). Sampling locations were situated inside and outside two Marine Protected Areas (MPAs) and each site encompassed one of the two typical coastal substrates of the Seychelles, granitic rocks or carbonate (coral) reefs. Ten sampling sites were inside protected areas and eight outside; six were located on granitic beds and twelve on coral ones (Table S1).

**Figure 1.**
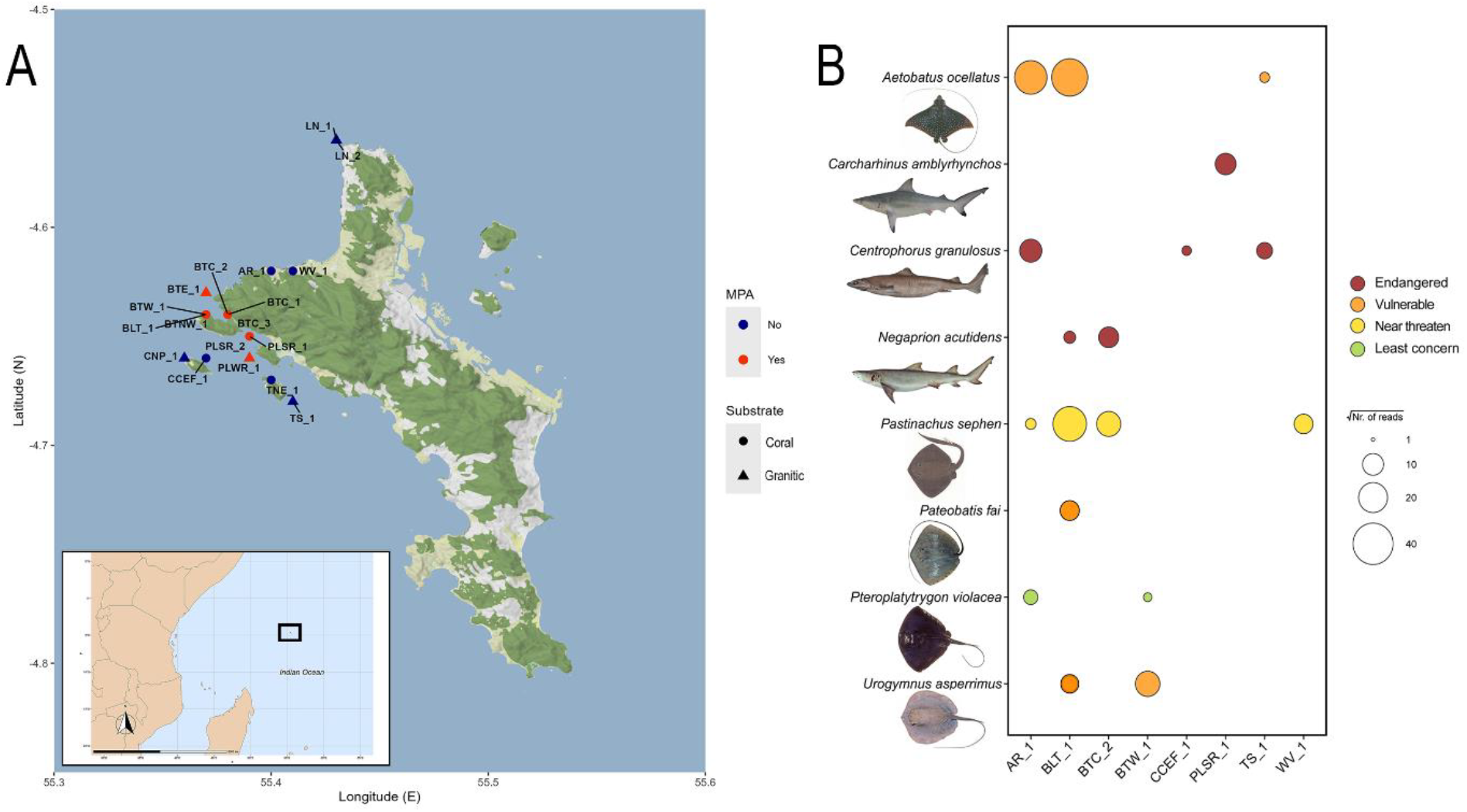
(A) Map of the 18 sampling sites along the coast of Mahé island on the Seychelles archipelago in the Indian Ocean. (B) Bubble plot representing the elasmobranch relative abundance per each sampling site; dot size correspond to the transformed (square-root) number of reads and colours are based on the IUCN conservation status of each species.

Environmental DNA samples were gathered through the passive soaking of sterile medical gauze rolls secured with zip-ties into a metaprobe (https://github.com/GiuliaMaiello/Metaprobe-2.1). Each metaprobe contained three rolls of gauze and was attached to the tank of recreational scuba divers who swam underwater in coastal waters. Dive times were kept virtually constant (44-46 minutes ranges). For each dive, we recorded the date, the location, geographic coordinates, the type of substrate, the maximum depth, the duration of the dive, the sea temperature, and whether a sample was taken in or outside marine protected areas (Table S1). At the end of each dive, metaprobes were detached from the diver’s tanks and the three gauze sub-samples were immediately retrieved and stored in separate 50 mL falcon tubes with silica gel grains to dry the gauze and preserve the DNA. Samples were frozen at -20 °C in the laboratory until subsequent processing.

### 2.2 Laboratory Procedures

DNA extraction and PCR preparation were carried out wearing full coverall suites, mask and hood, in a dedicated eDNA clean laboratory, where materials were sterilized using 10% bleach, rinsed with 70% ethanol and subsequently UV-irradiated, to minimize the risk of external contamination. The DNA was extracted directly from the gauze by cutting off small pieces from various parts of the rolls. The amount of gauze used for DNA extraction was adjusted to fit into a 1.5 mL Eppendorf tubes and the remaining gauze was stored in the original 50 mL tube at -20 °C. Extraction procedures followed a slightly modified version of the Purification of Total DNA from Animal Tissues protocol of the DNeasy Blood and Tissue Kit (Qiagen) (see details in the Supplementary methods). Three extraction negatives (one for each extraction day) were included to monitor the possibility and extent of contamination linked with extraction procedures and reagents.

The 57 DNA extractions (54 samples and three extraction controls) along with two PCR negative controls and one positive control (i.e., the DNA of the iridescent freshwater catfish *Pangasianodon hypophthalmus*) were PCR amplified targeting a ∼171 bp 12S ribosomal RNA fragment of the mitochondrial genome (Miya et al., 2015), using the Elas02 primers (forward: 5’-GTTGGTHAATCTCGTGCCAGC-3’; reverse: 5’-CATAGTAGGGTATCTAATCCTAGTTTG-3’), specifically designed to target elasmobranchs (Taberlet et al., 2018). The choice of this marker was driven by the necessity to capture the highest possible number of elasmobranch species – given their charismatic role in attracting diving tourism – while also ensuring an exhaustive coverage of teleost fauna, as recently been shown by Maiello et al. (2024), whose amplification and library preparation protocols were used (see details about PCR amplification and library preparation in the Supplementary method section).

The library was sequenced at 85 pM with 20% PhiX Control on an Illumina® iSeq™ 100 using the i1 Reagent v2 (300-cycle) (Illumina Inc.).

### 2.3 Bioinformatic Pipeline

Prior to bioinformatic processing of sequencing data, the quality of raw fastq files was assessed with FASTQC v 0.12.1 (Andrews, 2010). Forward and reverse reads were then merged using default settings of the function: ‘--fastq_mergepairs’ in VSEARCH v 2.23.0 (Rognes et al., 2016). Merged sequences were demultiplexed and assigned to samples based on their unique barcodes with CUATADAPT v 4.4 (Martin, 2011), allowing for a single mismatch (parameter: ‘-e’) in the barcode and primer region. We then filtered demultiplexed reads using the ‘--fastq_filter’ function in VSEARCH based on a maximum expected error of 1 (parameter: ‘--fastq_maxee’), a minimum length of 130 bp (parameter: ‘--fastq_minlen’), a maximum length of 210 bp (parameter: ‘--fastq_maxlen’), and without allowing the occurrence of ambiguous base calls (parameter: ‘--fastq_maxns 0’). Sequences passing the quality filtering thresholds were dereplicated using the ‘--derep_fulllength’ function in VSEARCH with default parameters. We removed chimeras with UCHIME (Edgar et al., 2011) and clustered the remaining sequences into Molecular Operational Taxonomical Units (MOTUs) with SWARM (Mahé et al., 2015) setting the threshold to d = 3. Finally, a frequency table was generated using the ‘--otutabout’ function in VSEARCH.

### 2.4 Reference Database and Taxonomic Assignment

Two reference databases for the Elas02 primer set were created using CRABS v 0.1.8 (Jeunen et al., 2023). The first one was generated by downloading all available 12S sequences from three online repositories (i.e., NCBI, EMBL and Mitofish), while the second database consisted solely of sequences from fish documented to exist in Seychelles waters, which were retrieved from the NCBI and Mitofish repositories. The full list of Seychelles fish (n = 1186) was obtained by selecting the ISO country code 690 in the RFISHBASE R package (Boettiger et al., 2012). For both reference databases, sequences were first downloaded from the multiple online repositories with the ‘db_download’ function and merged using the ‘db_merge’ function. Target amplicons were then extracted from sequences through in silico PCR analysis (‘insilico_pcr’ function) and pairwise global alignments (‘pga’ function). The curated databases were finally dereplicated (‘dereplicate’ function) and filtered (‘seq_cleanup’ function).

Taxonomic assignment was performed by using a variety of methods (1) the ‘-sintax’ function in USEARCH (Edgar, 2016) and (2) BLASTn (v2.11.0), using the Seychelles fish curated reference database; and (3) the ‘-sintax’ function against the general Elas02 12S database. The final taxonomic assignment of each MOTU was determined by seeking for a consensus among the three assignment methods, as detailed in the Supplementary methods.

Following taxonomic assignment, artefact sequences were filtered by merging reads based on taxon-dependent co-occurrence patterns as implemented in TOMBRAIDER v 0.1.0 (Jeunen et al., 2024), using default settings. The dataset was then filtered removing potential contamination noise, taking advantage of control samples (extraction and PCR negatives) using the MICRODECON v 1.0.2 R package (McKnight et al., 2019) with default settings (see Supplementary methods for further details on taxonomic assignment and final dataset refinement).

### 2.5 Statistical Analyses

All downstream statistical analyses were conducted in R v4.3.2 (R Core Team 2024). We first explored the abundance and spatial distribution of detected fish taxa, using square-root transformed reads as a semi-quantitative measure of abundance. A bubble plot was drawn to visualize the elasmobranch relative read count per each sampling site where sharks and rays were detected, highlighting the IUCN conservation status of each species. We also evaluated the abundance pattern of commercially valuable fish species, visualizing the transformed reads as a heatmap and testing differences in abundance patterns between MPA and non-MPA sites via a PERMANOVA test (Bray-Curtis distance), with the ‘adonis’ function in the VEGAN package (Oksanen et al., 2018). The commercial value of each species was assessed using information from Fishbase (Froese & Pauly, 2002), where the extent of species exploitation for human consumption is indicated, with the following six categories: highly commercial, commercial, minor commercial, subsistence fisheries, of potential interest, of no interest. Only taxa classified as ‘commercial’ and ‘highly commercial’ were included in the analysis of abundance patterns of commercially valuable species.

Boxplots were used to compare the alpha diversity (number of identified taxa) between MPA an non-MPA sites and between sampling location with carbonate (coral) or granitic rock substrate. The statistical significance of pair-wise comparisons was evaluated by Kruskal-Wallis test, following assessment of non-normality (Shapiro-Wilk test) and variance heterogeneity (Levene’s test). Differences in species assemblages between MPA/non-MPA and coral/granitic sites was visualised through a non-metric multidimensional scaling (nMDS) and tested via a PERMANOVA test, using the ‘metaMDS’ and the ‘adonis’ function in the VEGAN package, respectively. Distance matrices were calculated using Jaccard distance on a binary presence/absence dataset of all the taxa identified.

In addition to taxonomic diversity, the functional diversity of fish assemblages was also assessed, considering five functional traits: maximum size, depth range, aggregation behaviour, position in the water column and trophic category (Table S4). Trait values for each category were assigned to each taxon using information from Fishbase. Whenever the information about the trait characteristic of a species was missing, we took the trait value from the closest congeneric species exhibiting similar life history and ecology. The MFD package (Magneville et al., 2022) was then used to group taxa sharing the same trait values in unique functional entities, and boxplots were drawn to compare the mean number of functional entities between groups (MPA vs non-MPA and coral vs granitic). Functional redundancy (i.e., number of fish taxa divided by the number of functional entities) was calculated and the mean value estimates were compared between groups (Kruskal-Wallis test). Alluvial plots were drawn to visualize the relative proportion of trait values present in MPA/non-MPA and coral/granitic sites for each functional trait separately (i.e., maximum size, depth range, aggregation behaviour, position in the water column and trophic category). Both the number of taxonomic units and functional entities were contrasted with the sampling effort (number of samples), computing accumulation curves using the INEXT package (Hsieh et al., 2016). Accumulation curves were calculated and plotted separately for MPA/non-MPA and coral/granitic sites.

The MFD package was used also to compute functional diversity indices based on a multidimensional trait-space (‘alpha.fd.multidim’ function). First, pairwise trait-based distances for all sets of traits were calculated using Gower’s distance (Legendre, & Legendre, 1998) and then a principal coordinate analysis (PCoA) was performed on this distance matrix to generate the multidimensional space in which functional diversity indices were calculated. The mean absolute deviation index was used to assess the quality of the PCoA based multidimensional space and showed that the first four axes provided the best representation of the trait-based distances between taxa. Correlations between traits and axes of the functional space were computed and the index of functional richness was calculated in the 4D multidimensional space as the proportion of the functional space occupied by a certain assemblage (Villéger et al., 2008). Differences between functional richness estimates for MPA/non-MPA and coral/granitic sites were illustrated by projected convex hulls filled by the functional entities’ assemblages along axes of the functional space using the MFD package. Finally, the Jaccard coefficient was used to compute the functional dissimilarity between groups (MPA/non-MPA and coral/granitic), with the ‘beta.fd.multidim’ function of the MFD package, which parses the beta-diversity dissimilarity index into its additive components: ‘turnover’ (the replacement of entities between habitats) and ‘nestedness’ (the degree to which entities in one habitat are a smaller subset of the species found in the other).

## 3. Results

The Illumina iSeq run resulted in a total of 2.7 million raw reads. After bioinformatic analysis 962,312 reads were retained of which 219,977 were assigned to target taxa resulting in 174 unique fish species (Table S3), belonging to 127 different genera and 57 families (see Table S2 for further details on the reads removed at each main bioinformatic and filtering step). Three of the 18 total sampling sites (i.e., PLWR_1, LN_1 and BTNW_1) were removed from the final dataset because of their low read depth (< 1,000). Eight of the 174 identified taxa were elasmobranchs, and were detected in eight different sampling sites (Fig. 1B). Almost all of them are of conservation concern according to the IUCN red list: three species are considered endangered, three vulnerable, one near threatened and only one of least concern. Of the 166 teleosts, 78 were commercially important species and eight of them of particularly high commercial value (Table S3). Many of these commercially valuable species (71 out of 86, including *Cephalopholis argus, Chlorurus sordidus, Scarus niger, Rastrelliger kanagurta* and *Selar crumenophthalmus*) were more abundant (in term of square-root transformed reads) in the sites located inside Marine Protected Areas (Fig. 2), with differences in abundance patterns between MPA and non-MPA supported by PERMANOVA results (R^2^ = 0.11, p = 0.01).

**Figure 2.**
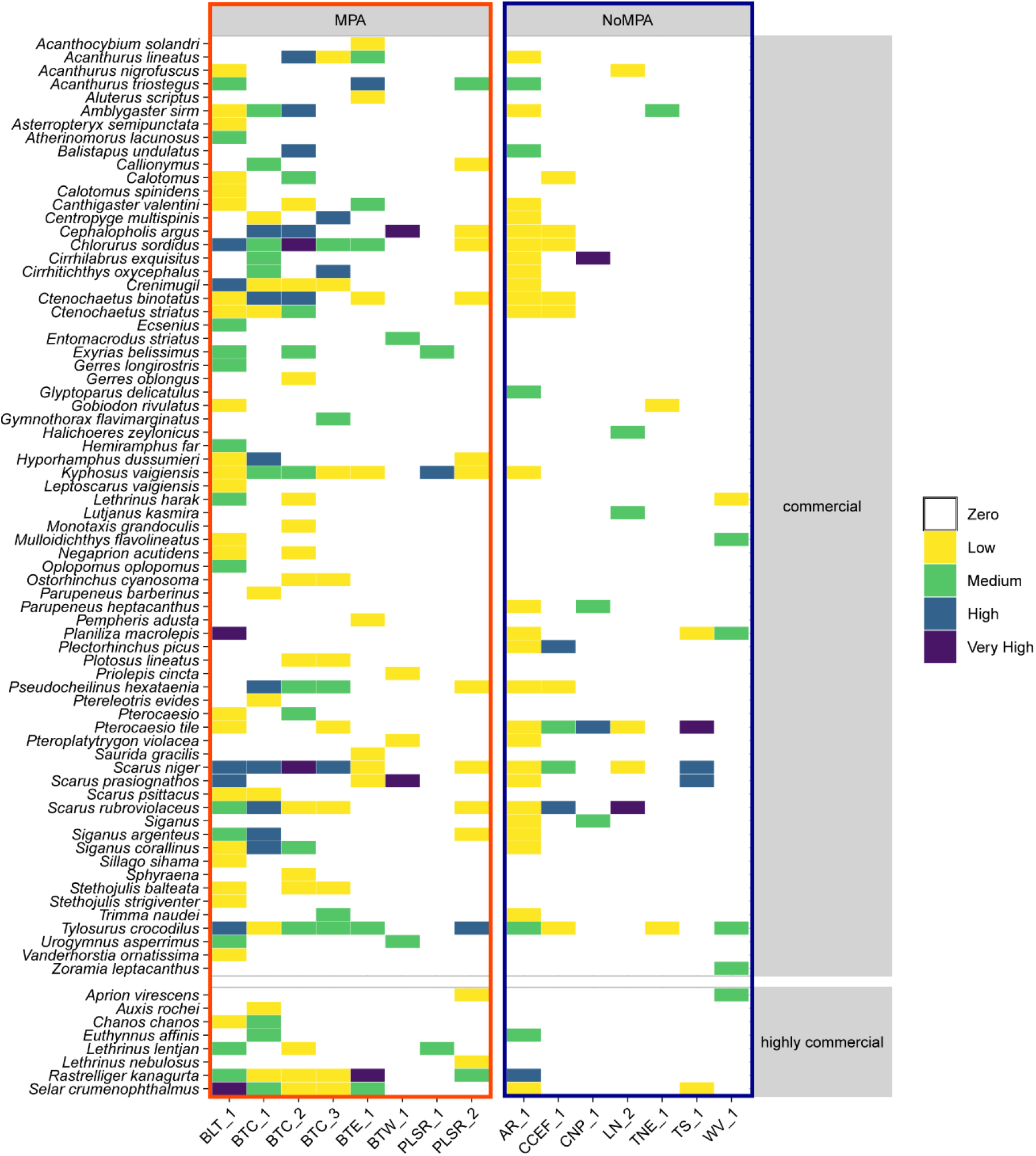
Quantitative composition of commercial and highly commercial teleosts in terms of transformed read counts for each sampling site, depicted by a heatmap representing the square-root of the number of reads. Sampling sites are separated between the ones located on MPA and non-MPA. Colours ranges are defined as follow: zero = 0, low = 0-10, medium = 10 – 20, high = 20 – 40, very high > 40.

The mean number of taxa was significantly higher in the MPA compared to the non-MPA sites and in the sampling locations with a carbonate substrate compared to the granitic ones (Fig. 3A and 3B; Kruskal–Wallis: H = 6.9, df = 1, p < 0.05 and H = 3.7, df = 1, p = 0.05, respectively). Despite the limited extent of the sampled area, the nMDS showed a fish community structure influenced by both the presence of MPAs and the type of substrate of sampling sites, with no significant association between protection status and type of substrate (Fisher’s exact test: odds ratio = 0.2, p = 0.05). Separation of MPA/non-MPA and coral/granitic sites was evident from Figure 3 (C and D) and was corroborated by the PERMANOVA results (Table S6).

**Figure 3.**
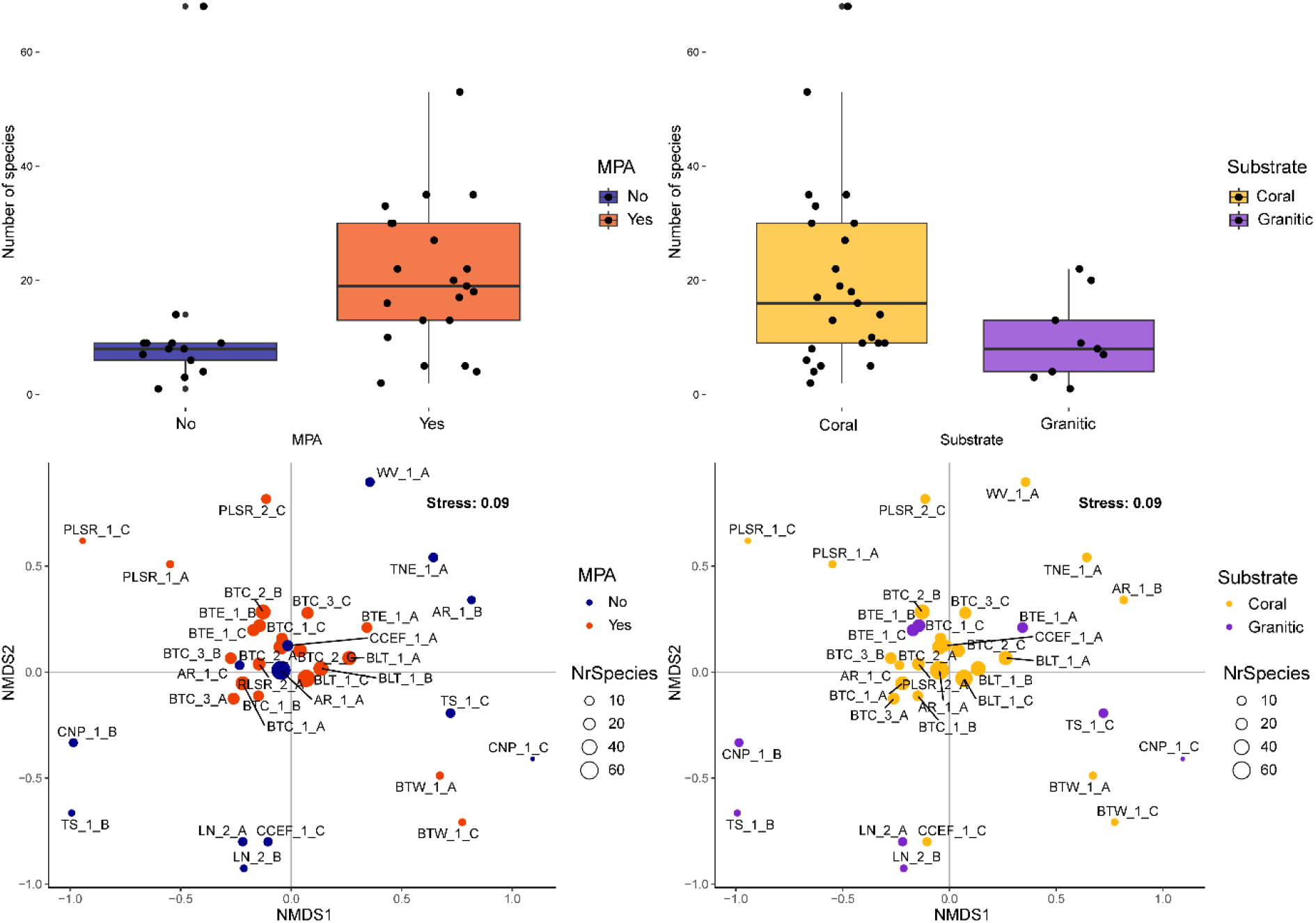
Comparison of sampling locations in terms of taxon detection; boxplots represent the number of species per sample detected in MPA/non-MPA (A) and coral/granitic sites (B). Pattern of the species assemblages across sampling sites, as returned by the non-metric multidimensional scaling (nMDS) with Jaccard distance. Sampling sites are coloured according to the presence or not of marine protected area (C), and the type of substrate (i.e., coral/granitic) (D).

A total of 112 unique functional entities were identified from the association of the 174 species with the five functional traits (Table S5). The accumulation curves attested that for all the considered groups (MPA/non-MPA, coral/granitic) the sampling effort was not sufficient to capture both the expected taxonomic and functional diversities (Fig. S1); however, the functional diversity curves were closer to the plateau.

Functional alpha and beta-diversity showed similar patterns with those observed for the taxonomic diversity. The mean number of functional entities was higher in the MPA compared to non-MPA sites and in the sampling locations with a carbonate substrate compared to the granitic ones (Fig. S2A and B), with no statistical differences in the functional redundancy among groups (mean values = 1.07 and 1.05 for MPA/non-MPA and H = 1.37, df = 1, p = 0.24; mean values = 1.07 and 1.03 for coral/granitic and H = 1.12, df = 1, p = 0.29). For each of the five functional trait (i.e., maximum size, depth range, aggregation behaviour, position in the water column and trophic category), the relative proportion of trait values (calculated as the proportion of functional entities exhibiting each trait value) was considerably higher in MPA and coral sites compared to non-MPA and granitic sites, as indicated by the greater stream thickness in the alluvial plot (Fig. S3). The nMDS based on trait data (Fig. S2C and D) showed a clear separation in two-dimensional space among MPA/non-MPA and coral/granitic sites, thereby strengthening the community patterns observed at the taxonomic level.

In the 4D multidimensional trait-space used to calculate the functional richness, the statistically significant traits driving PCoA axes are summarized in Table 1. The functional richness was slightly higher in the MPA sites compared to the ones located outside MPAs. Functional entities detected in MPA filled a functional surface corresponding to almost 90% of the global functional space, while the ones detected in non-MPA covered 76% of this surface (Fig. 4A and S4A), corresponding to a Jaccard functional dissimilarity of 0.28 (61% turnover and 39% nestedness components of beta-diversity). Interestingly, dissimilarities among surfaces were considerably larger when considering functional entities detected in coral and granitic sites, which filled a functional surface representing 99% and 45% of the total functional space, respectively. The Jaccard functional dissimilarity between coral and rock-dominated habitats was 0.54, with 98% of this value due to the nestedness component of beta diversity, highlighting that the taxa detected in rocky sites filled a subset of the coral functional space (Fig. 4B and S4B).

**Table 1.** Statistically significant traits driving PCoA axes in the 4D multidimensional trait-space used to calculate the functional richness. For each trait and axis statistical values (Kruskal-Wallis test) and p-values are given.

| Trait | Axis | Value | p.value |
| --- | --- | --- | --- |
| Size | PC3 | 0.825 | < 0.001 |
| Depth | PC1 | 0.066 | 0.017 |
| Depth | PC3 | 0.114 | 0.001 |
| Depth | PC4 | 0.591 | < 0.001 |
| Aggregation behaviour | PC2 | 0.86 | < 0.001 |
| Aggregation behaviour | PC3 | 0.126 | < 0.001 |
| Aggregation behaviour | PC4 | 0.044 | 0.034 |
| Water column position | PC1 | 0.157 | < 0.001 |
| Water column position | PC2 | 0.133 | < 0.001 |
| Water column position | PC3 | 0.142 | < 0.001 |
| Water column position | PC4 | 0.147 | < 0.001 |
| Trophic category | PC1 | 0.824 | < 0.001 |
| Trophic category | PC2 | 0.087 | 0.010 |
| Trophic category | PC4 | 0.095 | 0.007 |

**Figure 4.**
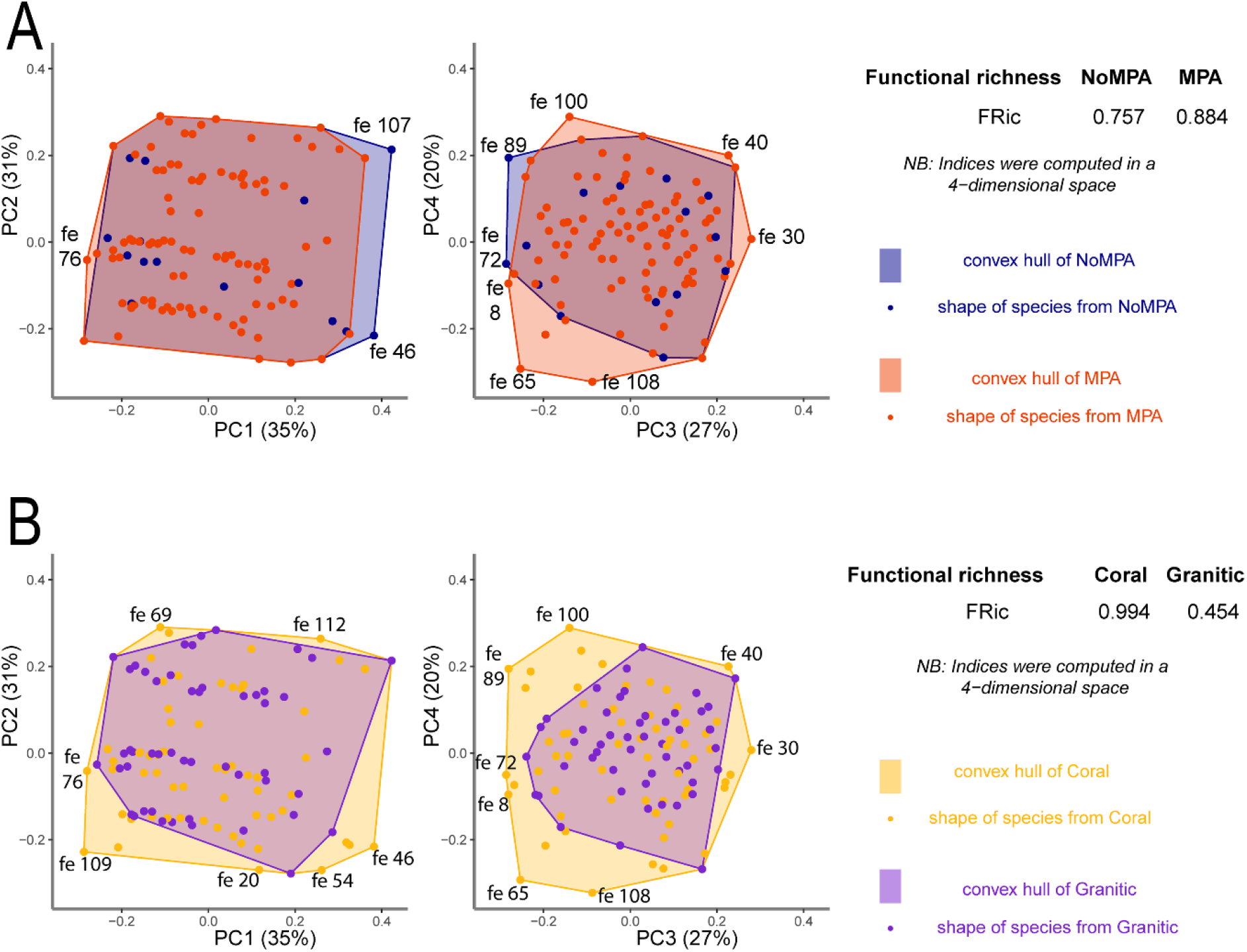
Comparison between functional richness estimates at the global scale, computed in a PCoA-based 4D functional space. Each subplot represents the functional space for a given pair of axes (PC1-PC2 and PC3-PC4). (A) Blue dots correspond to functional entities (fe) identified in non-MPA and orange dots to those detected in MPA. The blue polygon represents the trait space covered by fish entities detected in non-MPA, whereas the orange polygon represents the trait space covered by functional entities identified in MPA. (B) Yellow dots correspond to functional entities (fe) identified in coral sites and purple dots to those detected in the granitic ones. The yellow polygon represents the trait space covered by fish entities detected in coral sites, whereas the purple polygon represents the trait space covered by functional entities identified in the granitic ones. The functional entity (fe) numbers correspond to the ones in Table S5.

## 4. Discussion

The advent of eDNA-based biodiversity assessment is offering new solutions to the challenge of ecosystem monitoring under increasing climate and anthropogenic challenges. It is also becoming evident that standard water filtration protocols remain a major limitation for scaling up data collection (Hansen et al., 2020), especially in remote regions, with limited resources. Nimble low-tech passive methods, such as the metaprobe (Maiello et al., 2022) are amenable to potentially countless participatory science applications. Here we adopted it in association with the recreational activity of scuba divers enjoying the reefs on the west of Mahé island, in the Seychelles archipelago. Using an elasmobranch-specific 12S marker on eDNA metaprobe samples collected during 18 dives, we were able to identify 174 fish taxa. In line with previous evidence, we found that the Elas02 primers, although specifically designed to amplify elasmobranch DNA, also efficiently detect the typically more abundant teleost species (Maiello et al., 2024; Mariani et al., 2021), which represents a valuable emerging reality for biodiversity screening under strict budget constraints.

Seven of the elasmobranch species detected are considered of conservation concern according to the IUCN Red List. For instance, the grey reef shark *Carcharhinus amblyrhynchos* and the sicklefin lemon shark *Negaprion acutidens*, two very emblematic shark species of Indo-Pacific reefs, are listed as endangered. As other large-bodied sharks, they represent a major proportion of the high trophic-level predator biomass and play a crucial role in structuring reef communities (Speed et al., 2019). Their presence in three coral reef sites located in two Marine Protected Areas (i.e., Baie Ternay and Port Launay) suggests that these conservation measures may provide a lifeline for these severely threatened organisms (MacNeil et al., 2020). The other endangered shark, the gulper shark *Centrophorus granulosus*, is a deep-water species usually found between 200 and 600 metres, whose detection was not expected in shallow coastal waters. Despite several studies showing fine-scale spatial resolution and vertical stratification of marine environmental DNA data (Andruszkiewicz et al., 2017; Jeunen et al., 2019, 2020; Port et al., 2016), the possibility of DNA fragments dispersion from distant areas and depths cannot be ruled out. The presence of *C. granulosus* DNA in coastal waters could also come from residues of DNA in fishing nets; the gulper shark is known to be a common by-catch of demersal trawlers (Gray & Kennelly, 2018) and the sampling site where the species was more abundant (AR1 -Auberge Reef) is located on a highly populated area next to the small fishing harbour of Bel Ombre.

The 166 teleost species included the full breadth of iconic tropical reef taxa (Table S3), such as parrotfishes (e.g., *Calotomus spinidens, Chlorurus sordidus* and *Scarus spp*.), butterflyfishes of the genus *Chaetodon*, various wrasse species (e.g., *Cheilinus spp*., *Cirrhilabrus spp*., and *Hemigymnus spp*.) and surgeon fishes of the genera *Acanthurus* and *Ctenochaetus*. Of these, 78 species are considered of commercial importance for the fishing industry and eight of them are ‘highly commercial’. Interestingly, commercially valuable species appear invariably more abundant in the sampling sites located in marine protected areas, as evident from Figure 2 and statistically supported by PERMANOVA results (p = 0.02). For instance, five commercial species (i.e., *Cephalopholis argus, Chlorurus sordidus, Planiliza macrolepis, Scarus niger* and *Scarus prasiognathos*) and two highly commercial (i.e., the Indian mackerel *Rastrelliger kanagurta* and the Bigeye scad *Selar crumenophthalmus*) were particularly abundant (> 1,600 reads) in MPA sites. Coral reef and in general tropical fish populations are heavily impacted from intense fishing, which is leading to decline and even some local extinctions (Graham et al., 2011; Samoilys et al., 2022). An increase in abundance of intensely fished species in MPAs, where fishing is prohibited, lends further support around the effectiveness of these conservation measures. We found twice as many species on average in the sites located in Marine Protected Areas: (20.4 vs 11.9 between MPA and non-MPA respectively; H = 6.9, p < 0.05) (Fig. 3A). Coral habitats also returned around double the number of species than rocky ones (19.8 vs 9.7; H = 3.7, p = 0.05) (Fig. 3B). Interestingly, the sampling sites with the highest number of taxa were in the Baie Ternay Marine National Park (sites BLT and BTC) (Fig. 1A), which is a small protected patch (<1 km^2^) supporting a diverse array of marine habitats including mangroves, sea grass beds, rocky shores, and coral reefs reaching depths of nearly 40 meters. Given that more diverse fish communities are more resilient to the impacts of climate changes (Duffy et al., 2016), these results suggest that – if strategically placed – even very small protected areas may contribute to sustain diverse communities.

In line with recent studies that successfully used eDNA data to assess the functional diversity in various marine environments (Aglieri et al., 2021; Clay et al., 2024; Condachou et al., 2023; Roblet et al., 2024), including hyper-diverse coral reefs (Rey et al., 2023), here we identified 112 functional entities, collapsing the 174 species based on their values of the five considered functional traits. The reduction to functional diversity also allows to capture a greater proportion of the vast ecological diversity of coral reefs (Fig. S1), especially when sampling effort is a limiting factor. With increased participation of recreational scuba divers, it is possible to imagine a near future where increased sampling frequency and broader geographical cover can lead to comprehensive ecological assessment of the world’s most vulnerable coral reefs.

Alpha and beta diversity differences between MPA/non-MPA and coral/granitic sites were also reflected in the analyses at the functional level (Fig. S2 A-B). Functional diversity was higher for MPA and coral sites, indicating the presence of more heterogeneous trait combinations for functional entities exclusively associated with MPA and coral reef locations (Fig. 4 and Fig. S4). MPA sites were characterized by a wide range of functional trait values (Fig. S3), spanning from very small non-schooling bentho-demersal taxa such as gobies (e.g., *Asterropteryx semipunctata, Eviota guttata, Favonigobius reichei*), to medium-sized benthic species such as moray eels (*Gymnothorax melatremus* and *Gymnothorax flavimarginatus*). Interestingly, among the functional entities primarily detected in MPAs, there were also larger pelagic top predator species, such as the Blacktail reef shark (*Carcharhinus amblyrhynchos*), the snubnose dart (*Trachinotus blochi*), and the wahoo *(Acantocybium solandri*). High trophic-level predators are very important for their role as ecosystem engineers; top-predator decline has been associated to cascading effects in food webs, drastic modifications in their prey communities, and overall degradations of ecosystem functions and services (Estes et al., 2016; Heithaus et al., 2008; Myers et al., 2007). The role of large-bodied fishes in regulating biomass and associated ecosystem functions is such that their loss appears to be a major factor in overall biodiversity loss in tropical environments (Lefcheck et al., 2021).

Functional entities associated with coral sites had a substantially greater and more diverse range of functional values (Fig. S3). Almost all (98%) the dissimilarity with rocky sites was associated with the nestedness component of beta diversity, with the taxa detected in rocky sites merely constituting a subset of the coral functional space (Fig 4B and Fig. S4B). As expected, among the species uniquely characterizing coral locations we found very small, shallow, bentho-demersal species, which typically live and hide in crevices of coral reefs, such as blennies (e.g., *Enneapterygius tutuilae* and *Glyptoparus delicatulus*) and gobiids (e.g., *Vanderhorstia ornatissima, Pleurosicya mossambica, Eviota guttata*). There were also several pelagic or demersal schooler taxa, which are known to be preferentially detected by eDNA because of the large amount of DNA they release in the water column (Kelly et al., 2014; Maruyama et al., 2014). Among the schooling or facultative schoolers species, some were iconic species of coral reef assemblages (e.g., yellowfin surgeonfish *Acanthurus xanthopterus*, saddled puffer *Canthigaster valentini*, bluespotted grouper *Cephalopholis argus*).

Regardless of the modest geographic extent of the investigated area, the overall qualitative β-diversity distribution remarkably discriminated between the 15 sampling sites and reflected patterns of community structure influenced by both the presence of marine protected areas and the type of reef substrate. These differences in fish community composition between MPA/non-MPA and coral/rocky habitats involved both taxonomic and functional community elements, which aligns with growing evidence for the power of eDNA analysis to reveal and monitor key features of marine assemblages across habitats (Monuki et al., 2021; Port et al., 2016; Yamamoto et al., 2016).

What is perhaps most significant is that such evidence can now be gathered through simple, judicious, inexpensive agreements with citizen scientists – in this case recreational scuba divers – hence by-passing the often substantial financial and logistic challenges associated with bespoke marine surveys. An added value to this framework is the empowering of the environmentally-aware public to actively participate in the stewarding of the ecosystems that underpin their passion. Although there is a requirement for further method refinement and more efforts towards the establishment of frequent, coordinated, and comprehensive sampling programmes, an avenue is now paved towards an affordable, engaging, inclusive, universal approach to directly surveying and stewarding virtually every coastal sea.

## Acknowledgments

We thank David Rowe and Gabriella Thomas for their precious support during diving and sampling procedures and their involvement and contributions to this project.

The fieldwork for this study was conducted as part of the research project entitled ‘Enhancing marine resource stewarding in the Seychelles using environmental DNA’ approved by the Seychelles Bureau of Standard and the Seychelles Parks & Gardens Authority and led by Victor Burgos Rubio in collaboration with Global Vision International (GVI).

The study was supported by the UK Natural Environment Research Council [“SpongeDNA” grant NE/T007028/1].

## Data availability

Raw sequencing data and metadata will be made available in a Dryad repository upon acceptance of this manuscript. Bioinformatic pipelines, sample barcodes, final datasets and R scripts used for statistical analysis and to generate figures will be publicly available from https://github.com/GiuliaMaiello/ Opportunistic-diver-assisted-eDNA-sampling-Seychelles.

