## Supplementary material for "Opportunistic diver-assisted eDNA sampling unpicks fine-scale ecological and conservation signals in tropical reef fishes"

### Supplementary Methods

#### DNA extraction:

The pieces of gauze were incubated at 56°C in a thermomixer overnight (~16 hours) with 720 µl Buffer ATL and 80 µl Proteinase K (20 µg/ml). DNA was then extracted according to the manufacturer protocol, with reagent volumes adjusted based on the initial volume of ATL Buffer. Total DNA was finally concentrated through a DNeasy Mini spin column and eluted in 100 µl of elution buffer. The elution through the spin column was repeated twice to increase the final DNA concentration in the eluate.

#### PCR amplification and library preparation:

Primer pairs were uniquely tagged for each sample with 8 bp oligo-tags to unequivocally identify samples during demultiplexing and reduce the risk of cross-contamination and/or tag switching during Illumina sequencing. Each tag differed by at least three base pairs from all other tags and was preceded by 2–4 degenerate bases (Ns) to improve sequence diversity. We prepared PCR reactions in a total volume of 20 µl for each sample containing 10 µl MyFi™ Mix (Meridian Bioscience), 0.16 µl of Bovine Serum Albumin (20 mg/ml, Thermo Scientific), 5.84 µl of UltraPure™ Distilled Water (Invitrogen), 1 µl of each forward and reverse primer (10 µM, Eurofins), and 2 µl of template DNA. All PCRs were run in triplicates under the following thermocycling conditions: polymerase activation at 94°C for 15 mins, followed by 40 cycles of 94°C for 1 min, 54°C for 1 min, 72°C for 1 min, and finishing with a final elongation at 72°C for 5 min followed by a 4°C hold.

PCR triplicates were then pooled and visualized through gel electrophoresis on a 2% agarose gel (150 ml 1X TBE buffer with 3 g agarose powder) stained with 1.5 µl SYBRsafe dye (Invitrogen) to ensure the successful amplification of target fragments. PCR products were purified using 1:1 ratio of PCR product to 30 µl of Mag-Bind® Total Pure NGS magnetic beads (Omega Bio-Tek) following the manufacturer's protocol (Bronner et al., 2009).

Purified DNA was quantified using a Qubit Flex™ 4.0 fluorometer with the Qubit™ dsDNA HS Assay Kit (Invitrogen) and the DNA quantification of all samples was used to normalize and pool samples in equimolar concentration for library preparation. The fragment length of pooled samples was analysed with an Agilent 4200 TapeStation and High Sensitivity D1000 ScreenTape (Agilent Technologies) which showed the presence of a single peak of the expected size. We thus performed end- repair, adapter ligation and library PCR amplification using the NEXTFLEX® Rapid DNA-Seq Kit 2.0 for Illumina® platforms (PerkinElmer) according to the manufacturer's protocol. After the ligation of Illumina adapters, the library was checked again on the Tape Station that showed secondary products (e.g., adaptor dimers) remained, which were removed by an additional 1x ratio magnetic bead clean-up. The cleaned library was quantified by quantitative PCR (qPCR) on a Rotor-Gene Q (Qiagen) using the NEBNext® Library Quant Kit for Illumina® (New England Biolabs) and then diluted to 1 nM according to qPCR concentrations. The diluted library was re-quantified along with a PhiX Control using qPCR before sequencing

#### Taxonomic assignment:

Taxonomic assignment was performed by using three methods (1) the '-sintax' function in USEARCH (Edgar, 2016) and (2) BLASTn (v2.11.0), using the Seychelles fish curated reference database; and (3) the '-sintax' function against the general Elas02 12S database. The consensus among the three assignment methods for the final taxonomic assignment of each MOTU was determined as follow. First, MOTUs for which all three methods provided the same taxonomic classification were assigned accordingly. When consensus among all three methods was not possible, MOTUs that were assigned to the same taxon by both Sintax and BLAST with the Seychelles fish species reference database were assigned based on this agreement. Finally, remaining MOTUs were assigned according to Sintax 12S general reference database results.

#### Dataset final refinement:

Further refinement of the data set consisted of removing: (1) non-marine taxa (e.g., human) and non-target fish species (i.e., fish species not expected in Seychelles marine waters whose detection could not result from DNA shed by natural populations in the study area, but rather from human-mediated contamination, such as the Atlantic salmon *Salmo salar* and the European perch *Perca fluviatilis*), (2) taxa that could not unambiguously be assigned to the genus or species rank, (3) MOTUs showing < 90% sequence identity, (4) samples not reaching a total abundance of 1,000 reads. Finally, to reduce the effects of low-abundance false positives due to tag switching and/or cross-contamination, only detections with an occurrence > 2 within each sample were retained for downstream ecological analyses (Schnell & Bohmann, 2015).

**Table S1 -** Name, date, location, geographical coordinates (datum=WGS84), substrate, protection status (i.e., MPA/non MPA), depth, duration of the dive and sea temperature are given for each sampling site.

| Site | Date | Location | Lat (S) | Long (E) | Subst | MPA | Depth (m) | Duration (min) | Sea Temp (°C) |
|---|---|---|---|---|---|---|---|---|---|
| AR_1 | 08/03/23 | Auberge Reef | 4.62 | 55.4 | Carb | No | 5.8 | 44 | 30 |
| BLT_1 | 16/06/23 | Baie Ternay Lighthouse | 4.64 | 55.37 | Carb | Yes | 7 | 44 | 30 |
| BTC_1 | 16/06/23 | Baie Ternay Reef Centre | 4.64 | 55.38 | Carb | Yes | 8.2 | 44 | 30 |
| BTC_2 | 16/06/23 | Baie Ternay Reef Centre | 4.64 | 55.38 | Carb | Yes | 8 | 46 | 30 |
| BTC_3 | 16/06/23 | Baie Ternay Reef Centre | 4.64 | 55.38 | Carb | Yes | 7.6 | 45 | 30 |
| BTE_1 | 09/03/23 | Baie Ternay Reef North East | 4.63 | 55.37 | Gran | Yes | 14 | 45 | 30 |
| BTNW_1 | 14/03/23 | Baie Ternay Reef North West | 4.64 | 55.37 | Carb | Yes | 9.2 | 45 | 30 |
| BTW_1 | 08/03/23 | Baie Ternay Reef West | 4.64 | 55.37 | Carb | Yes | 5.6 | 44 | 30 |
| CCEF_1 | 14/03/23 | Conception Central East Face | 4.66 | 55.37 | Carb | No | 10.5 | 46 | 30 |
| CNP_1 | 14/03/23 | Conception North Point | 4.66 | 55.36 | Gran | No | 13.8 | 46 | 29 |
| LN_1 | 03/04/23 | L'ilot North Face | 4.56 | 55.43 | Gran | No | 16 | 45 | 30 |
| LN_2 | 03/04/23 | L'ilot North Face | 4.56 | 55.43 | Gran | No | 7 | 45 | 30 |
| PLSR_1 | 20/03/23 | Port Launay South Reef | 4.65 | 5.39 | Carb | Yes | 10 | 45 | 30 |
| PLSR_2 | 20/03/23 | Port Launay South Reef | 4.65 | 5.39 | Carb | Yes | 5.5 | 45 | 30 |
| PLWR_1 | 13/03/23 | Port Launay West Rocks | 4.66 | 5.39 | Gran | Yes | 9.7 | 44 | 30 |
| TNE_1 | 15/03/23 | Therese North East | 4.67 | 5.4 | Carb | No | 12.8 | 45 | 30 |
| TS_1 | 09/03/23 | Therese South | 4.68 | 55.41 | Gran | No | 16.4 | 44 | 30 |
| WV_1 | 20/03/23 | White Villa Reef | 4.62 | 55.41 | Carb | No | 7.5 | 45 | 30 |

**Table S2** – Total number of reads and MOTUs at different steps of the bioinformatic and filtering process.

| | Reads | Unique MOTUs |
|---|---|---|
| **Total raw reads** | 2,700,00 | - |
| **After bioinformatic analysis** | 962,312 | 837 |
| **Assigned taxonomy (Metazoa)**<br>**Unassigned** | 898,960<br>63,352 | 632<br>205 |
| **Assigned to target taxa** | 440,210 | 582 |
| **Assigned to non-target taxa (e.g., human)** | 443,940 | 24 |
| **Assigned to non-target fish (i.e., not recorded in the Seychelles)** | 14,810 | 26 |
| **Target taxa after filtering** | 219,977 | 174 |

**Table S3 -** List of all the fish taxa (i.e., genus and species) detected. Information about the corresponding class, IUCN conservation status and importance for the fishing industry are provided for each species.

| Scientific name | Class | IUCN | Fisheries |
|---|---|---|---|
| *Abudefduf* | Actinopterygii | Least concern | minor commercial |
| *Abudefduf septemfasciatus* | Actinopterygii | Least concern | minor commercial |
| *Acanthocybium solandri* | Actinopterygii | Least concern | commercial |
| *Acanthurus* | Actinopterygii | Least concern | minor commercial |
| *Acanthurus leucosternon* | Actinopterygii | Least concern | minor commercial |
| *Acanthurus lineatus* | Actinopterygii | Least concern | commercial |
| *Acanthurus nigrofuscus* | Actinopterygii | Least concern | commercial |
| *Acanthurus triostegus* | Actinopterygii | Least concern | commercial |
| *Acanthurus xanthopterus* | Actinopterygii | Least concern | minor commercial |
| *Aethaloperca rogaa* | Actinopterygii | Least concern | minor commercial |
| *Aetobatus ocellatus* | Chondrichthyes | Vulnerable | non commercial |
| *Albula glossodonta* | Actinopterygii | Vulnerable | minor commercial |
| *Aluterus scriptus* | Actinopterygii | Least concern | commercial |
| *Amblyeleotris diagonalis* | Actinopterygii | Least concern | non commercial |
| *Amblygaster sirm* | Actinopterygii | Least concern | commercial |
| *Amblyglyphidodon indicus* | Actinopterygii | Least concern | non commercial |
| *Anampses caeruleopunctatus* | Actinopterygii | Least concern | minor commercial |
| *Anampses meleagrides* | Actinopterygii | Least concern | minor commercial |
| *Anyperodon leucogrammicus* | Actinopterygii | Least concern | minor commercial |
| *Apolemichthys* | Actinopterygii | Least concern | minor commercial |
| *Aprion virescens* | Actinopterygii | Least concern | highly commercial |
| *Arctozenus risso* | Actinopterygii | Least concern | non commercial |
| *Arothron* | Actinopterygii | Least concern | minor commercial |
| *Arothron mappa* | Actinopterygii | Least concern | minor commercial |
| *Asterropteryx semipunctata* | Actinopterygii | Least concern | commercial |
| *Atherinomorus lacunosus* | Actinopterygii | Least concern | commercial |
| *Auxis rochei* | Actinopterygii | Least concern | highly commercial |
| *Azurina lepidolepis* | Actinopterygii | Least concern | non commercial |
| *Balistapus undulatus* | Actinopterygii | Least concern | commercial |
| *Caesio lunaris* | Actinopterygii | Least concern | minor commercial |
| *Caesio teres* | Actinopterygii | Least concern | minor commercial |
| *Caesio varilineata* | Actinopterygii | Least concern | minor commercial |
| *Callionymus* | Actinopterygii | Least concern | commercial |
| *Callogobius* | Actinopterygii | Least concern | non commercial |
| *Callogobius maculipinnis* | Actinopterygii | Least concern | non commercial |
| *Calotomus* | Actinopterygii | Least concern | commercial |
| *Calotomus spinidens* | Actinopterygii | Least concern | commercial |
| *Canthigaster valentini* | Actinopterygii | Least concern | commercial |
| *Caranx papuensis* | Actinopterygii | Least concern | minor commercial |
| *Carcharhinus amblyrhynchos* | Chondrichthyes | Endangered | minor commercial |

| | | | |
|---|---|---|---|
| *Centrophorus granulosus* | Chondrichthyes | Endangered | minor commercial |
| *Centropyge multispinis* | Actinopterygii | Least concern | commercial |
| *Cephalopholis argus* | Actinopterygii | Least concern | commercial |
| *Cephalopholis sexmaculata* | Actinopterygii | Least concern | subsistence fisheries |
| *Chaetodon auriga* | Actinopterygii | Least concern | minor commercial |
| *Chaetodon kleinii* | Actinopterygii | Least concern | subsistence fisheries |
| *Chaetodon trifasciatus* | Actinopterygii | Least concern | minor commercial |
| *Chanos chanos* | Actinopterygii | Least concern | highly commercial |
| *Cheilinus* | Actinopterygii | Least concern | minor commercial |
| *Cheilio inermis* | Actinopterygii | Least concern | minor commercial |
| *Cheilodipterus quinquelineatus* | Actinopterygii | Least concern | minor commercial |
| *Cheilopogon* | Actinopterygii | Least concern | non commercial |
| *Chlorurus sordidus* | Actinopterygii | Least concern | commercial |
| *Choerodon* | Actinopterygii | Not Evaluated | non commercial |
| *Chromis opercularis* | Actinopterygii | Least concern | non commercial |
| *Chromis weberi* | Actinopterygii | Least concern | non commercial |
| *Cirrhilabrus exquisitus* | Actinopterygii | Data deficient | commercial |
| *Cirrhitichthys oxycephalus* | Actinopterygii | Least concern | commercial |
| *Cirripectes* | Actinopterygii | Least concern | non commercial |
| *Coris formosa* | Actinopterygii | Least concern | minor commercial |
| *Crenimugil* | Actinopterygii | Least concern | commercial |
| *Ctenochaetus binotatus* | Actinopterygii | Least concern | commercial |
| *Ctenochaetus striatus* | Actinopterygii | Least concern | commercial |
| *Diaphus splendidus* | Actinopterygii | Least concern | non commercial |
| *Echidna nebulosa* | Actinopterygii | Least concern | minor commercial |
| *Echidna polyzona* | Actinopterygii | Least concern | subsistence fisheries |
| *Ecsenius* | Actinopterygii | Least concern | commercial |
| *Enneapterygius* | Actinopterygii | Least concern | non commercial |
| *Enneapterygius tutuilae* | Actinopterygii | Least concern | non commercial |
| *Entomacrodus striatus* | Actinopterygii | Least concern | commercial |
| *Euthynnus affinis* | Actinopterygii | Least concern | highly commercial |
| *Eviota* | Actinopterygii | Least concern | non commercial |
| *Eviota guttata* | Actinopterygii | Least concern | non commercial |
| *Eviota shimadai* | Actinopterygii | Least concern | non commercial |
| *Exyrias belissimus* | Actinopterygii | Least concern | commercial |
| *Favonigobius reichei* | Actinopterygii | Least concern | non commercial |
| *Fistularia commersonii* | Actinopterygii | Least concern | minor commercial |
| *Gerres longirostris* | Actinopterygii | Least concern | commercial |
| *Gerres oblongus* | Actinopterygii | Least concern | commercial |
| *Glyptoparus delicatulus* | Actinopterygii | Least concern | commercial |
| *Gnathanodon speciosus* | Actinopterygii | Least concern | minor commercial |
| *Gobiodon rivulatus* | Actinopterygii | Least concern | commercial |
| *Gomphosus caeruleus* | Actinopterygii | Least concern | minor commercial |
| *Gymnomuraena zebra* | Actinopterygii | Least concern | minor commercial |
| *Gymnothorax flavimarginatus* | Actinopterygii | Least concern | commercial |
| *Gymnothorax javanicus* | Actinopterygii | Least concern | subsistence fisheries |

| | | | |
|---|---|---|---|
| *Gymnothorax melatremus* | Actinopterygii | Least concern | minor commercial |
| *Gymnothorax pindae* | Actinopterygii | Least concern | minor commercial |
| *Halichoeres* | Actinopterygii | Least concern | minor commercial |
| *Halichoeres hortulanus* | Actinopterygii | Least concern | minor commercial |
| *Halichoeres zeylonicus* | Actinopterygii | Least concern | commercial |
| *Helcogramma* | Actinopterygii | Least concern | non commercial |
| *Hemigymnus fasciatus* | Actinopterygii | Least concern | minor commercial |
| *Hemigymnus melapterus* | Actinopterygii | Least concern | minor commercial |
| *Hemiramphus far* | Actinopterygii | Not Evaluated | commercial |
| *Hirundichthys speculiger* | Actinopterygii | Least concern | minor commercial |
| *Hypoatherina temminckii* | Actinopterygii | Not Evaluated | subsistence fisheries |
| *Hyporhamphus dussumieri* | Actinopterygii | Not Evaluated | commercial |
| *Iniistius pentadactylus* | Actinopterygii | Least concern | minor commercial |
| *Kyphosus cinerascens* | Actinopterygii | Least concern | minor commercial |
| *Kyphosus vaigiensis* | Actinopterygii | Least concern | commercial |
| *Leptoscarus vaigiensis* | Actinopterygii | Least concern | commercial |
| *Lethrinus harak* | Actinopterygii | Least concern | commercial |
| *Lethrinus lentjan* | Actinopterygii | Least concern | highly commercial |
| *Lethrinus nebulosus* | Actinopterygii | Least concern | highly commercial |
| *Lethrinus obsoletus* | Actinopterygii | Least concern | minor commercial |
| *Limnichthys nitidus* | Actinopterygii | Least concern | non commercial |
| *Lutjanus kasmira* | Actinopterygii | Least concern | commercial |
| *Monotaxis grandoculis* | Actinopterygii | Least concern | commercial |
| *Mulloidichthys flavolineatus* | Actinopterygii | Least concern | commercial |
| *Myripristis* | Actinopterygii | Least concern | minor commercial |
| *Myripristis kuntee* | Actinopterygii | Least concern | minor commercial |
| *Negaprion acutidens* | Chondrichthyes | Endangered | commercial |
| *Neoniphon argenteus* | Actinopterygii | Least concern | non commercial |
| *Neoniphon sammara* | Actinopterygii | Least concern | minor commercial |
| *Novaculichthys taeniourus* | Actinopterygii | Least concern | minor commercial |
| *Oplopomus oplopomus* | Actinopterygii | Least concern | commercial |
| *Ostorhinchus cyanosoma* | Actinopterygii | Least concern | commercial |
| *Ostorhinchus holotaenia* | Actinopterygii | Least concern | non commercial |
| *Ostracion cubicum* | Actinopterygii | Not Evaluated | minor commercial |
| *Oxycheilinus digramma* | Actinopterygii | Least concern | minor commercial |
| *Parapriacanthus ransonneti* | Actinopterygii | Not Evaluated | non commercial |
| *Parupeneus barberinus* | Actinopterygii | Least concern | commercial |
| *Parupeneus heptacanthus* | Actinopterygii | Least concern | commercial |
| *Pastinachus sephen* | Chondrichthyes | Near Threatened | minor commercial |
| *Pateobatis fai* | Chondrichthyes | Vulnerable | minor commercial |
| *Pempheris adusta* | Actinopterygii | Not Evaluated | commercial |
| *Planiliza macrolepis* | Actinopterygii | Least concern | commercial |
| *Platax orbicularis* | Actinopterygii | Least concern | minor commercial |
| *Plectorhinchus picus* | Actinopterygii | Not Evaluated | commercial |
| *Pleurosicya mossambica* | Actinopterygii | Least concern | non commercial |
| *Plotosus lineatus* | Actinopterygii | Not Evaluated | commercial |

| | | | |
|---|---|---|---|
| *Pomacanthus imperator* | Actinopterygii | Least concern | minor commercial |
| *Pomacentrus caeruleus* | Actinopterygii | Least concern | non commercial |
| *Pomacentrus trilineatus* | Actinopterygii | Least concern | non commercial |
| *Priolepis cincta* | Actinopterygii | Least concern | commercial |
| *Pseudocheilinus hexataenia* | Actinopterygii | Least concern | commercial |
| *Ptereleotris evides* | Actinopterygii | Least concern | commercial |
| *Pterocaesio* | Actinopterygii | Least concern | commercial |
| *Pterocaesio tile* | Actinopterygii | Least concern | commercial |
| *Pteroplatytrygon violacea* | Chondrichthyes | Least concern | commercial |
| *Pygoplites diacanthus* | Actinopterygii | Least concern | minor commercial |
| *Rastrelliger kanagurta* | Actinopterygii | Least concern | highly commercial |
| *Rhabdamia gracilis* | Actinopterygii | Not Evaluated | non commercial |
| *Sargocentron* | Actinopterygii | Least concern | minor commercial |
| *Saurida gracilis* | Actinopterygii | Least concern | commercial |
| *Scarus niger* | Actinopterygii | Least concern | commercial |
| *Scarus prasiognathos* | Actinopterygii | Least concern | commercial |
| *Scarus psittacus* | Actinopterygii | Least concern | commercial |
| *Scarus rubroviolaceus* | Actinopterygii | Least concern | commercial |
| *Schindleria praematura* | Actinopterygii | Least concern | non commercial |
| *Scolopsis* | Actinopterygii | Least concern | subsistence fisheries |
| *Selar crumenophthalmus* | Actinopterygii | Least concern | highly commercial |
| *Siganus* | Actinopterygii | Least concern | commercial |
| *Siganus argenteus* | Actinopterygii | Least concern | commercial |
| *Siganus corallinus* | Actinopterygii | Least concern | commercial |
| *Siganus rivulatus* | Actinopterygii | Least concern | minor commercial |
| *Sillago sihama* | Actinopterygii | Least concern | commercial |
| *Sphyraena* | Actinopterygii | Not Evaluated | commercial |
| *Spratelloides delicatulus* | Actinopterygii | Least concern | minor commercial |
| *Stegastes lacrymatus* | Actinopterygii | Not Evaluated | non commercial |
| *Stethojulis balteata* | Actinopterygii | Least concern | commercial |
| *Stethojulis strigiventer* | Actinopterygii | Least concern | commercial |
| *Sufflamen chrysopterum* | Actinopterygii | Least concern | minor commercial |
| *Synodus* | Actinopterygii | Least concern | minor commercial |
| *Taeniamia fucata* | Actinopterygii | Least concern | non commercial |
| *Trachinotus baillonii* | Actinopterygii | Least concern | minor commercial |
| *Trachinotus blochii* | Actinopterygii | Least concern | minor commercial |
| *Trimma naudei* | Actinopterygii | Least concern | commercial |
| *Tylosurus crocodilus* | Actinopterygii | Least concern | commercial |
| *Urogymnus asperrimus* | Chondrichthyes | Vulnerable | commercial |
| *Vanderhorstia ornatissima* | Actinopterygii | Least concern | commercial |
| *Verulux cypselurus* | Actinopterygii | Least concern | non commercial |
| *Zoramia leptacanthus* | Actinopterygii | Least concern | commercial |

**Table S4 -** Functional traits and their relative categories and values considered in the present study.

| Functional trait | Trait categories | Trait values |
|---|---|---|
| **Maximum length (Size)** | Very small | < 10 cm |
| | Small | 10-30 cm |
| | Medium-size | 30-50 cm |
| | Large | 50-100cm |
| | Very large | > 100 cm |
| **Depth** | Shallow | < 20 m |
| | Medium-depth | Up to 50 m |
| | Deep | Up to 200 m |
| | Very deep | Up to more than 200 m |
| **Aggregation behaviour** | Non-schooling | Solitary |
| | Facultative schooler | Can form schools |
| | Schooler | Always in schools |
| **Water column position** | Benthic | Live on the bottom |
| | Demersal | Live in contact with the bottom |
| | Pelagic | Live in the water column |
| **Trophic category** | Herbivore | Algae and phytoplankton |
| | Omnivore I | Preference for vegetables |
| | Omnivore II | Preference for animals |
| | Carnivore | Preference for invertebrates |
| | Piscivore | Preference for fish |

**Table S5** – List of all the functional entities identified from the association of the 174 species with functional traits. For each functional entity the scientific name of all the species and the trait value of each functional trait (i.e., size, depth, aggregation behaviour, water column position and trophic category) are given.

| Functional entities | Scientific name | Size | Depth | Aggregation behaviour | Water column position | Trophic category |
|---|---|---|---|---|---|---|
| fe_1 | *Anampses meleagrides*<br>*Apolemichthys* sp.<br>*Cirrhitichthys oxycephalus*<br>*Halichoeres* sp.<br>*Halichoeres hortulanus*<br>*Halichoeres zeylonicus*<br>*Pseudocheilinus hexataenia* | small | medium-depth | non-schooling | demersal | Carnivore |
| fe_2 | *Cirrhilabrus exquisitus*<br>*Neoniphon argenteus*<br>*Pempheris adusta*<br>*Scolopsis* sp.<br>*Stethojulis balteata*<br>*Stethojulis strigiventer* | small | shallow | schooler | demersal | Carnivore |
| fe_3 | *Caesio lunaris*<br>*Caesio teres*<br>*Caesio varilineata*<br>*Pterocaesio* sp.<br>*Pterocaesio tile* | medium-sized | medium-depth | schooler | pelagic | Carnivore |
| fe_4 | *Callogobius* sp.<br>*Callogobius maculipinnis*<br>*Enneapterygius* sp.<br>*Enneapterygius tutuilae*<br>*Helcogramma* sp. | very small | medium-depth | non-schooling | demersal | Carnivore |
| fe_5 | *Anampses caeruleopunctatus*<br>*Gomphosus caeruleus*<br>*Oxycheilinus digramma*<br>*Sufflamen chrysopterum* | medium-sized | medium-depth | non-schooling | demersal | Carnivore |
| fe_6 | *Cheilodipterus quinquelineatus*<br>*Chromis weberi*<br>*Ecsenius* sp.<br>*Sargocentron* sp. | small | medium-depth | facultative schooler | demersal | Carnivore |
| fe_7 | *Calotomus* sp.<br>*Calotomus spinidens* | medium-sized | deep | facultative schooler | demersal | Herbivore |
| fe_8 | *Asterropteryx semipunctata*<br>*Eviota guttata*<br>*Vanderhorstia ornatissima* | very small | shallow | non-schooling | benthic | Carnivore |
| fe_9 | *Cheilio inermis*<br>*Lethrinus obsoletus*<br>*Monotaxis grandoculis* | large | medium-depth | facultative schooler | demersal | Carnivore |
| fe_10 | *Chromis opercularis*<br>*Iniistius pentadactylus*<br>*Taeniamia fucata* | small | medium-depth | schooler | demersal | Carnivore |
| fe_11 | *Eviota* sp.<br>*Eviota shimadai*<br>*Priolepis cincta* | very small | medium-depth | non-schooling | benthic | Carnivore |
| fe_12 | *Gerres longirostris*<br>*Hemigymnus fasciatus* | medium-sized | medium-depth | facultative schooler | demersal | Carnivore |

| | | | | | | |
|---|---|---|---|---|---|---|
| | *Hemigymnus melapterus* | | | | | |
| fe_13 | *Negaprion acutidens*<br>*Pastinachus sephen*<br>*Pateobatis fai* | very large | deep | non-schooling | demersal | Piscivore |
| fe_14 | *Abudefduf* sp.<br>*Abudefduf septemfasciatus* | small | shallow | schooler | demersal | Omnivore II |
| fe_15 | *Acanthurus* sp.<br>*Acanthurus leucosternon*<br>*Scarus rubroviolaceus* | large | medium-depth | facultative schooler | demersal | Herbivore |
| fe_16 | *Acanthurus nigrofuscus*<br>*Siganus rivulatus* | small | medium-depth | schooler | demersal | Herbivore |
| fe_17 | *Aethaloperca rogaa*<br>*Cephalopholis sexmaculata* | large | deep | facultative schooler | demersal | Piscivore |
| fe_18 | *Albula glossodonta*<br>*Lethrinus harak* | large | shallow | facultative schooler | demersal | Carnivore |
| fe_19 | *Amblyeleotris diagonalis*<br>*Exyrias belissimus* | small | medium-depth | non-schooling | benthic | Carnivore |
| fe_20 | *Aprion virescens*<br>*Fistularia commersonii* | very large | deep | facultative schooler | demersal | Piscivore |
| fe_21 | *Arothron* sp.<br>*Arothron mappa* | large | medium-depth | non-schooling | demersal | Omnivore II |
| fe_22 | *Atherinomorus lacunosus*<br>*Parapriacanthus ransonneti* | small | medium-depth | schooler | pelagic | Carnivore |
| fe_23 | *Azurina lepidolepis*<br>*Ostorhinchus holotaenia* | very small | medium-depth | schooler | demersal | Carnivore |
| fe_24 | *Callionymus* sp.<br>*Choerodon* sp. | small | deep | non-schooling | demersal | Carnivore |
| fe_25 | *Chlorurus sordidus*<br>*Siganus corallinus* | medium-sized | medium-depth | facultative schooler | demersal | Herbivore |
| fe_26 | *Coris formosa*<br>*Plectorhinchus picus* | large | medium-depth | non-schooling | demersal | Carnivore |
| fe_27 | *Gymnothorax melatremus*<br>*Gymnothorax pindae* | medium-sized | medium-depth | non-schooling | benthic | Piscivore |
| fe_28 | *Hirundichthys speculiger*<br>*Hypoatherina temminckii* | small | shallow | schooler | pelagic | Carnivore |
| fe_29 | *Kyphosus cinerascens*<br>*Kyphosus vaigiensis* | large | medium-depth | schooler | demersal | Omnivore I |
| fe_30 | *Lethrinus nebulosus*<br>*Parupeneus barberinus* | large | deep | facultative schooler | demersal | Carnivore |
| fe_31 | *Mulloidichthys flavolineatus*<br>*Parupeneus heptacanthus* | medium-sized | deep | facultative schooler | demersal | Carnivore |
| fe_32 | *Myripristis* sp.<br>*Myripristis kuntee* | small | deep | schooler | demersal | Omnivore II |
| fe_33 | *Oplopomus oplopomus*<br>*Ptereleotris evides* | small | shallow | non-schooling | demersal | Carnivore |
| fe_34 | *Ostorhinchus cyanosoma*<br>*Zoramia leptacanthus* | very small | shallow | schooler | demersal | Carnivore |
| fe_35 | *Scarus niger*<br>*Scarus psittacus* | medium-sized | medium-depth | non-schooling | demersal | Herbivore |
| fe_36 | *Siganus* sp.<br>*Siganus argenteus* | medium-sized | medium-depth | schooler | demersal | Herbivore |
| fe_37 | *Acanthocybium solandri* | very large | shallow | non-schooling | pelagic | Piscivore |
| fe_38 | *Acanthurus lineatus* | medium-sized | shallow | facultative schooler | demersal | Omnivore I |

| | | | | | | |
|---|---|---|---|---|---|---|
| fe_39 | *Acanthurus triostegus* | small | deep | facultative schooler | demersal | Herbivore |
| fe_40 | *Acanthurus xanthopterus* | large | deep | schooler | demersal | Omnivore I |
| fe_41 | *Aetobatus ocellatus* | very large | medium-depth | facultative schooler | demersal | Carnivore |
| fe_42 | *Aluterus scriptus* | very large | shallow | non-schooling | demersal | Omnivore I |
| fe_43 | *Amblygaster sirm* | small | deep | schooler | pelagic | Carnivore |
| fe_44 | *Amblyglyphidodon indicus* | very small | shallow | facultative schooler | demersal | Carnivore |
| fe_45 | *Anyperodon leucogrammicus* | large | medium-depth | non-schooling | demersal | Piscivore |
| fe_46 | *Arctozenus risso* | medium-sized | very deep | facultative schooler | pelagic | Piscivore |
| fe_47 | *Auxis rochei* | large | medium-depth | schooler | pelagic | Piscivore |
| fe_48 | *Balistapus undulatus* | medium-sized | medium-depth | non-schooling | demersal | Omnivore II |
| fe_49 | *Canthigaster valentini* | small | medium-depth | schooler | demersal | Omnivore II |
| fe_50 | *Caranx papuensis* | large | medium-depth | facultative schooler | pelagic | Piscivore |
| fe_51 | *Carcharhinus amblyrhynchos* | very large | very deep | facultative schooler | pelagic | Piscivore |
| fe_52 | *Centrophorus granulosus* | very large | very deep | non-schooling | demersal | Piscivore |
| fe_53 | *Centropyge multispinis* | small | medium-depth | facultative schooler | demersal | Herbivore |
| fe_54 | *Cephalopholis argus* | large | shallow | facultative schooler | demersal | Piscivore |
| fe_55 | *Chaetodon auriga* | small | deep | facultative schooler | demersal | Omnivore II |
| fe_56 | *Chaetodon kleinii* | small | deep | non-schooling | demersal | Omnivore II |
| fe_57 | *Chaetodon trifasciatus* | small | medium-depth | non-schooling | demersal | Omnivore II |
| fe_58 | *Chanos chanos* | very large | shallow | schooler | demersal | Omnivore I |
| fe_59 | *Cheilinus* sp. | very large | deep | non-schooling | demersal | Carnivore |
| fe_60 | *Cheilopogon* sp. | medium-sized | shallow | non-schooling | pelagic | Carnivore |
| fe_61 | *Cirripectes* sp. | very small | medium-depth | non-schooling | benthic | Omnivore I |
| fe_62 | *Crenimugil* sp. | large | shallow | schooler | demersal | Herbivore |
| fe_63 | *Ctenochaetus binotatus* | small | medium-depth | non-schooling | demersal | Herbivore |
| fe_64 | *Ctenochaetus striatus* | small | medium-depth | facultative schooler | demersal | Omnivore I |
| fe_65 | *Diaphus splendidus* | very small | very deep | non-schooling | pelagic | Carnivore |
| fe_66 | *Echidna nebulosa* | very large | shallow | non-schooling | benthic | Carnivore |
| fe_67 | *Echidna polyzona* | large | shallow | non-schooling | benthic | Carnivore |
| fe_68 | *Entomacrodus striatus* | small | shallow | non-schooling | demersal | Herbivore |

| | | | | | | |
|---|---|---|---|---|---|---|
| fe_69 | *Euthynnus affinis* | very large | deep | schooler | pelagic | Carnivore |
| fe_70 | *Favonigobius reichei* | very small | shallow | facultative schooler | demersal | Omnivore II |
| fe_71 | *Gerres oblongus* | medium-sized | medium-depth | schooler | demersal | Carnivore |
| fe_72 | *Glyptoparus delicatulus* | very small | shallow | non-schooling | benthic | Herbivore |
| fe_73 | *Gnathanodon speciosus* | very large | deep | schooler | demersal | Carnivore |
| fe_74 | *Gobiodon rivulatus* | very small | deep | non-schooling | demersal | Carnivore |
| fe_75 | *Gymnomuraena zebra* | very large | medium-depth | non-schooling | benthic | Carnivore |
| fe_76 | *Gymnothorax flavimarginatus* | very large | deep | non-schooling | benthic | Carnivore |
| fe_77 | *Gymnothorax javanicus* | very large | medium-depth | non-schooling | benthic | Piscivore |
| fe_78 | *Hemiramphus far* | medium-sized | shallow | schooler | pelagic | Herbivore |
| fe_79 | *Hyporhamphus dussumieri* | medium-sized | shallow | schooler | pelagic | Omnivore II |
| fe_80 | *Leptoscarus vaigiensis* | medium-sized | shallow | facultative schooler | demersal | Herbivore |
| fe_81 | *Lethrinus lentjan* | large | deep | non-schooling | demersal | Carnivore |
| fe_82 | *Limnichthys nitidus* | very small | shallow | schooler | demersal | Omnivore II |
| fe_83 | *Lutjanus kasmira* | medium-sized | deep | schooler | demersal | Carnivore |
| fe_84 | *Neoniphon sammara* | medium-sized | medium-depth | schooler | demersal | Piscivore |
| fe_85 | *Novaculichthys taeniourus* | medium-sized | shallow | non-schooling | demersal | Carnivore |
| fe_86 | *Ostracion cubicum* | medium-sized | medium-depth | non-schooling | demersal | Omnivore I |
| fe_87 | *Planiliza macrolepis* | large | shallow | schooler | demersal | Omnivore I |
| fe_88 | *Platax orbicularis* | large | medium-depth | facultative schooler | demersal | Omnivore I |
| fe_89 | *Pleurosicya mossambica* | very small | medium-depth | non-schooling | benthic | Omnivore II |
| fe_90 | *Plotosus lineatus* | medium-sized | medium-depth | facultative schooler | benthic | Carnivore |
| fe_91 | *Pomacanthus imperator* | medium-sized | deep | non-schooling | demersal | Carnivore |
| fe_92 | *Pomacentrus caeruleus* | small | shallow | non-schooling | demersal | Omnivore II |
| fe_93 | *Pomacentrus trilineatus* | small | shallow | non-schooling | demersal | Omnivore I |
| fe_94 | *Pteroplatytrygon violacea* | large | deep | non-schooling | pelagic | Carnivore |
| fe_95 | *Pygoplites diacanthus* | small | deep | facultative schooler | demersal | Carnivore |
| fe_96 | *Rastrelliger kanagurta* | medium-sized | deep | schooler | pelagic | Omnivore II |
| fe_97 | *Rhabdamia gracilis* | very small | deep | schooler | demersal | Carnivore |

| | | | | | | |
|---|---|---|---|---|---|---|
| fe_98 | *Saurida gracilis* | medium-sized | shallow | non-schooling | benthic | Piscivore |
| fe_99 | *Scarus prasiognathos* | large | medium-depth | schooler | demersal | Herbivore |
| fe_100 | *Schindleria praematura* | very small | deep | schooler | demersal | Omnivore II |
| fe_101 | *Selar crumenophthalmus* | large | shallow | schooler | pelagic | Carnivore |
| fe_102 | *Sillago sihama* | medium-sized | shallow | schooler | demersal | Carnivore |
| fe_103 | *Sphyraena* sp. | very large | medium-depth | non-schooling | pelagic | Piscivore |
| fe_104 | *Spratelloides delicatulus* | very small | medium-depth | schooler | pelagic | Omnivore II |
| fe_105 | *Stegastes lacrymatus* | small | medium-depth | non-schooling | demersal | Omnivore I |
| fe_106 | *Synodus* sp. | small | shallow | non-schooling | benthic | Piscivore |
| fe_107 | *Trachinotus baillonii* | large | shallow | schooler | pelagic | Piscivore |
| fe_108 | *Trachinotus blochii* | very large | shallow | facultative schooler | pelagic | Carnivore |
| fe_109 | *Trimma naudei* | very small | medium-depth | facultative schooler | benthic | Carnivore |
| fe_110 | *Tylosurus crocodilus* | very large | shallow | facultative schooler | pelagic | Piscivore |
| fe_111 | *Urogymnus asperrimus* | very large | very deep | non-schooling | demersal | Carnivore |
| fe_112 | *Verulux cypselurus* | very small | shallow | schooler | pelagic | Omnivore II |

**Table S6** – Results of PERMANOVA (Permutational Multivariate Analysis of Variance) analyses carried out to explore: taxa composition changes in relation to (A) protection status (i.e., MPA/non MPA) and (B) substrate (i.e., coral/granitic) of sampling sites; functional entities composition changes in relation to (C) protection status (i.e., MPA/non MPA) and (D) substrate (i.e., coral/granitic) of sampling sites. Values are related to Jaccard coefficients.

| | | Df | **SumsOfSqs** | **F.Model** | **$R^2$** | **p-value** |
|---|---|---|---|---|---|---|
| **A** | **MPA** | 1 | 0.72 | 1.99 | 0.06 | 0.004 |
| | **Residuals** | 32 | 11.64 | | 0.94 | |
| | **Total** | 33 | 12.36 | | 1.00 | |
| **B** | **Substrate** | 1 | 0.58 | 1.32 | 0.05 | 0.02 |
| | **Residuals** | 32 | 11.78 | | 0.95 | |
| | **Total** | 33 | 12.36 | | 1.00 | |
| **C** | **MPA** | 1 | 0.81 | 2.13 | 0.06 | < 0.001 |
| | **Residuals** | 32 | 12.22 | | 0.94 | |
| | **Total** | 33 | 13.04 | | 1.00 | |
| **D** | **Substrate** | 1 | 0.53 | 1.36 | 0.04 | 0.05 |
| | **Residuals** | 32 | 12.50 | | 0.96 | |
| | **Total** | 33 | 13.03 | | 1.00 | |

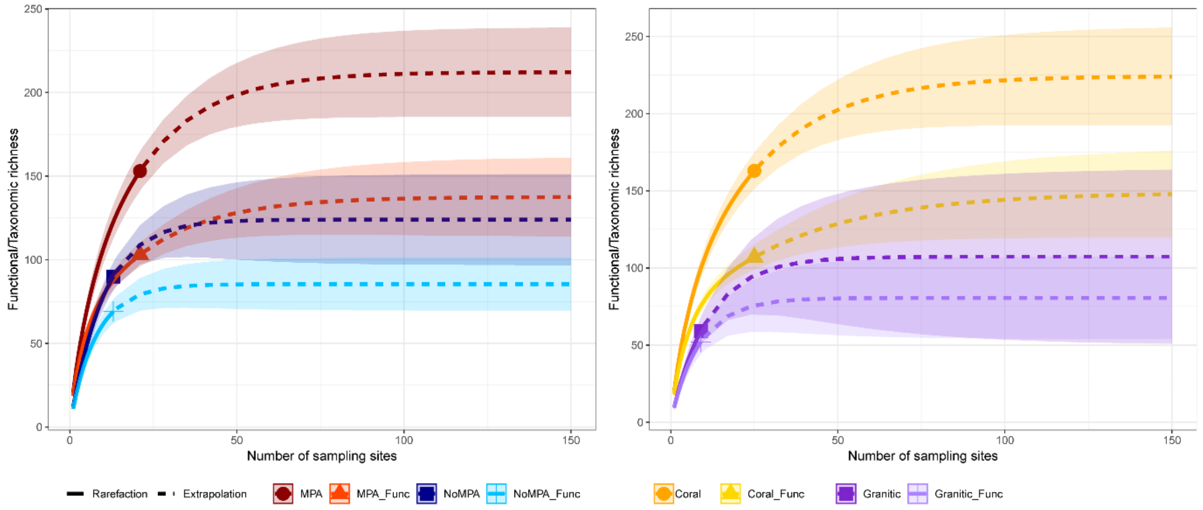

**Figure S1 -** Rarefaction curves depicting the total taxa and functional entities detected in the collected samples. Curves are split based on where the samples were collected: MPA/non MPA (A) and granitic/coral substrate (B).

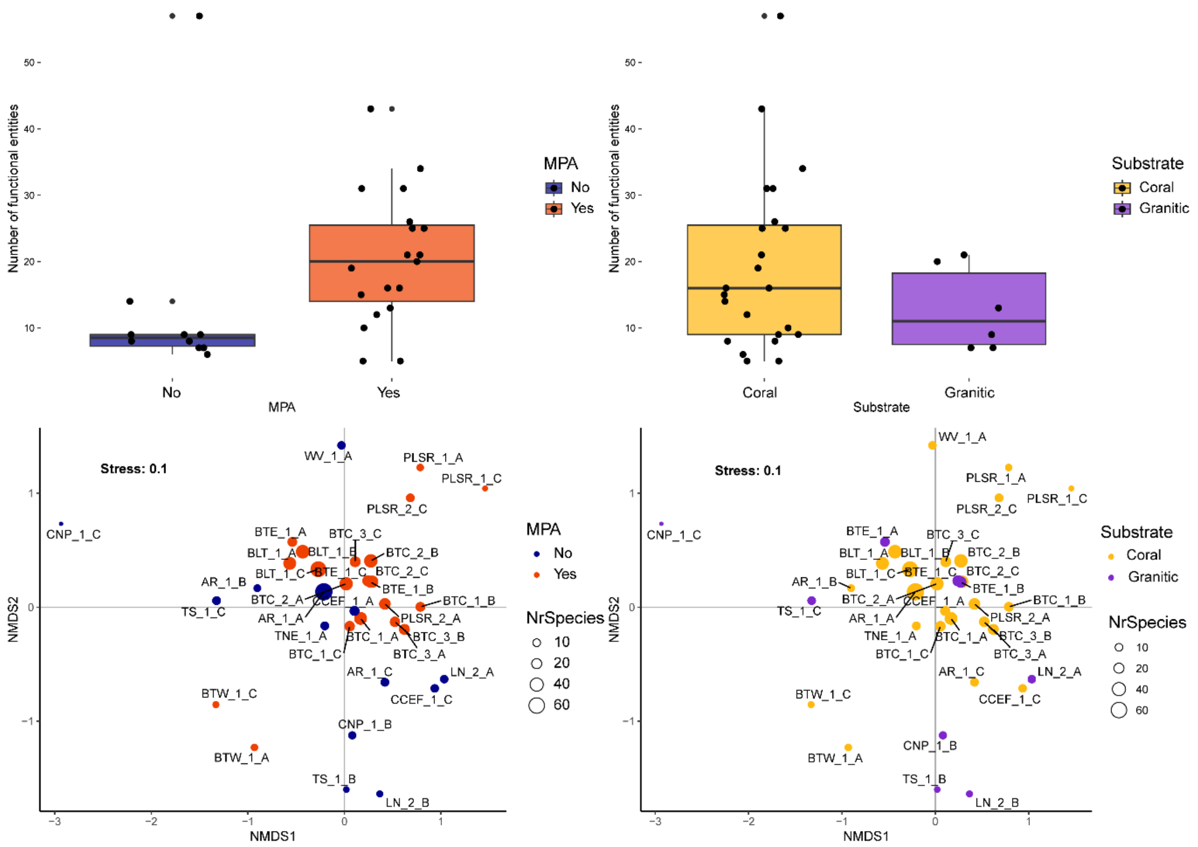

**Figure S2** - Comparison of sampling locations in terms of functional entity detection; boxplots represent the number of functional entities per sample detected in MPA/non MPA (A) and coral/granitic sites (B). Pattern of the functional entity assemblages across sampling sites, as returned by the non-metric multidimensional scaling (nMDS) with Jaccard distance. Sampling sites are coloured according to the presence or not of marine protected area (C), and the type of substrate (i.e., coral/granitic) (D).

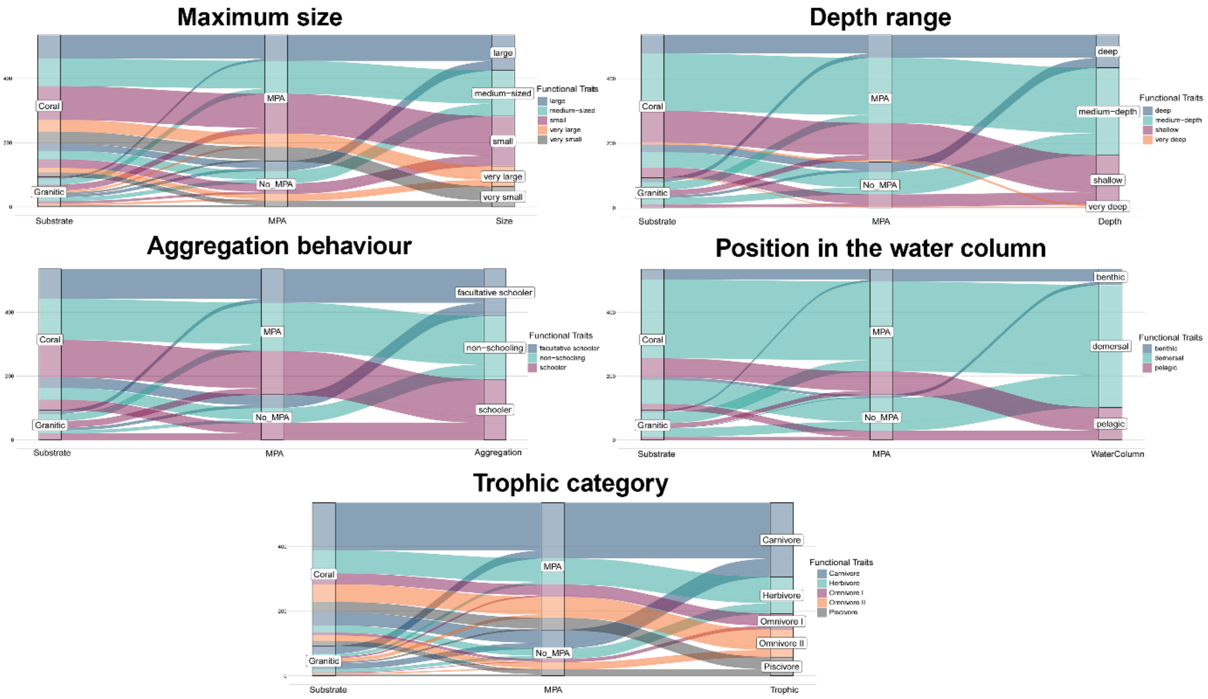

**Figure S3 –** Relative proportion of trait values present in MPA/non MPA and Coral/Granitic sites, depicted as an alluvial plot for each functional trait separately (i.e., maximum size, depth range, aggregation behaviour, position in the water column and trophic category). Stream thickness indicates the proportion of functional entities exhibiting each trait value.

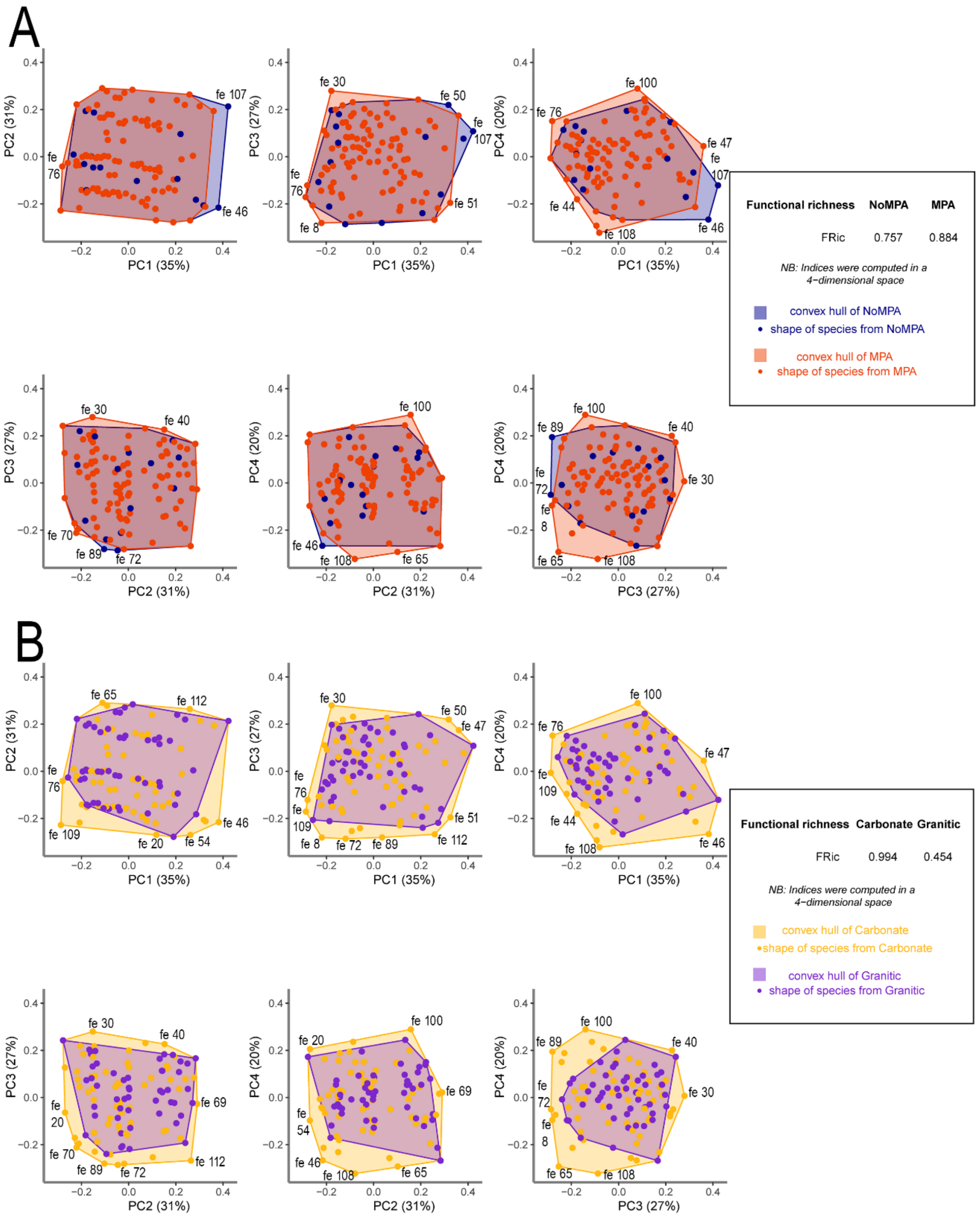

**Figure S4** – Comparison between functional richness estimates at the global scale, computed in a PCoA-based 4D functional space. Each subplot represents the functional space for a given pair of axes. (A) Blue dots correspond to functional entities identified in non-MPA and orange dots to those detected in MPA. The blue polygon represents the trait space covered by fish entities detected in non-MPA, whereas the orange polygon represents the trait space covered by functional entities identified in MPA. (B) Yellow dots correspond to functional entities identified in coral sites and purple dots to those detected in the granitic ones. The yellow polygon represents the trait space covered by fish entities detected in coral sites, whereas the purple polygon represents the trait space covered by functional entities identified in the granitic ones. The functional entity (fe) numbers correspond to the ones in Table S5.
